# Drug Screening with Zebrafish Visual Behavior Identifies Carvedilol as a Potential Treatment for Retinitis Pigmentosa

**DOI:** 10.1101/2020.07.28.225789

**Authors:** Logan Ganzen, MeeJung Ko, Mengrui Zhang, Rui Xie, Yongkai Chen, Liyun Zhang, Rebecca James, Jeff Mumm, Richard van Rijn, Wenxuan Zhong, Chi Pui Pang, Mingzhi Zhang, Motokazu Tsujikawa, Yuk Fai Leung

## Abstract

Retinitis Pigmentosa (RP) is an incurable inherited retinal degeneration affecting approximately 1 in 4,000 individuals globally. The goal of this work was to identify drugs that can help patients suffering from the disease. To accomplish this, we screened drugs on a zebrafish RP model. This model expresses a truncated human *rhodopsin* transgene (*Tg(rho:Hsa*.*RH1_Q344X*)) causing significant rod degeneration by 7 days post-fertilization (dpf). Consequently, the larvae displayed a deficit in visual motor response (VMR) under scotopic condition. The diminished VMR was leveraged to screen an ENZO SCREEN-WELL® REDOX library since oxidative stress is postulated to play a role in RP progression. Our screening identified a beta-blocker, carvedilol, that ameliorated the deficient VMR of the RP larvae and increased their rod number. Carvedilol can act directly on rods as it affected the adrenergic pathway in a rod-like human Y79 cell line. Since carvedilol is an FDA-approved drug, our findings suggest that carvedilol can potentially be repurposed to treat RP patients.

**Summary Statement:** This paper presents the utilization of zebrafish visual behavior, a novel paradigm to screen and identify drugs to treat retinitis pigmentosa, an incurable retinal-degenerative disease.

## Introduction

Retinitis Pigmentosa (RP) is an incurable retinal-degenerative disease affecting approximately 1 in 4,000 individuals globally (Hamel, 2006; Hartong et al., 2006; O’Neal and Luther, 2019). It is caused by multiple mutations found in at least 65 causative genes with different modes of inheritance (RetNet database: <https://sph.uth.edu/RetNet/>) (Daiger et al., 2013; Daiger et al., 2014; Hartong et al., 2006; Sorrentino et al., 2016). Patients suffering from RP have a cost burden of over $7,000 per year on average higher than healthy individuals (Frick et al., 2012). When patients lose their vision, their quality of life is lower and may suffer from increased likelihood of injury, and increased anxiety and depression (Welp et al., 2016). Unfortunately, there are currently no effective treatment options available for patients suffering from the disease. Research into technologies such as gene therapies, stem-cell therapy, and retinal prosthesis is being performed, however these options are still experimental and costly (Ganzen et al., 2017). This highlights the urgent need for RP therapeutics that are effective and inexpensive. To address this need, we developed a platform to perform phenotypic drug screening with a zebrafish RP model.

The zebrafish can provide a powerful system to model RP, and they have been used to model a number of human retinal-degenerative diseases (Brockerhoff and Fadool, 2011; Gross and Perkins, 2008; Link and Collery, 2015; Morris, 2011; Nakao et al., 2012). These models include transgenic zebrafish expressing human *rhodopsin* (*RHO*) with autosomal dominant mutations found in RP patients (Y. Sasamoto et al., 2010). Up to 30 percent of RP cases are autosomal dominant, and of all autosomal dominant cases, and approximately 30 percent arise due to over 150 mutations in *RHO* (Athanasiou et al., 2018; Daiger et al., 2014; Rossmiller et al., 2012). These mutations include Q344X/Q344ter, a truncation mutation, which shortens *RHO* (Sung et al., 1991). Q344X RHO lacks a VXPX ciliary trafficking motif leading to its mislocalization to the inner segment and apoptotic cell death (Concepcion and Chen, 2010; Portera-Cailliau et al., 1994; Sung et al., 1994). Patients with this mutation suffer an early onset, severe form of autosomal-dominant RP (Jacobson et al., 1991; Kremmer et al., 1997; Sung et al., 1994). In zebrafish, a transgenic model was made to express a human Q344X *RHO* in rods under zebrafish *rho* promoter (Nakao et al., 2012). This model exhibits significant rod degeneration as early as 5 days post-fertilization (dpf). The model also possesses a nose EGFP reporter in the transgenic cassette, which allows for efficient mutant screening starting at 2 dpf. Previous work with the Q344X zebrafish has shown that adenylyl cyclase (ADCY) inhibition can lead to modest rod survival (Nakao et al., 2012). It is hypothesized that aberrant ADCY signaling was activated by the mislocalized RHO in the inner segment, which would ultimately trigger apoptosis (Alfinito and Townes-Anderson, 2002; Nakao et al., 2012). However, it has also been shown that the activation of mislocalized RHO is not necessary to induce cell death (Tam et al., 2006). These findings indicate that Q344X can cause rod degeneration through more than one mechanism. In this study, we utilized this Q344X zebrafish model to develop an *in vivo* drug-screening platform for RP.

The zebrafish is an ideal model for *in vivo* screening for drugs to treat RP (Ganzen et al., 2017) due to its low cost of use, high fecundity, amenability to genetic manipulation (Patton and Zon, 2001). It can also facilitate RP drug discovery because of the rapid development of its visual system (Fadool and Dowling, 2008). In particular, zebrafish rod precursors begin to differentiate into rods as early as 36 hours post-fertilization (hpf) in the ventral region of the retina by expressing *rho* (Dowling, 2012; Hensley et al., 2011; Morris and Fadool, 2005). The rod outer segments begin to form by 50 hpf, and fully formed outer segments have been found as early as 4 dpf (Branchek and Bremiller, 1984; Fadool, 2003; Schmitt and Dowling, 1999). These rods begin to form synapses by 62 hpf (Schmitt and Dowling, 1994; Schmitt and Dowling, 1999). The earliest visually-evoked startle can be detected by 68 hpf (Easter, Jr. and Nicola, 1996). After that, several visual behaviors gradually appear from 3 dpf to 5 dpf, including the optokinetic response and the visual motor response (VMR) (Chhetri et al., 2014; Easter and Nicola, 1997; Emran et al., 2007; Liu et al., 2015). The VMR is a startle response triggered by a sudden light onset or offset, which results in increased locomotor behavior (Emran et al., 2007; Emran et al., 2008; Ganzen et al., 2017; Gao et al., 2014; Gao et al., 2016; Liu et al., 2015). This behavior can be measured from multiple larvae simultaneously in 96-well plate format and is thus ideal for high-throughput, *in vivo* drug screening experiments (Deeti et al., 2014; Ganzen et al., 2017). The VMR has been utilized to identify oculotoxic drugs, and discover drugs that can benefit retinal degeneration (Deeti et al., 2014; Zhang et al., 2016). Zebrafish have also been used to perform high-throughput drug screening based on fluorescent signals in the retina, but this approach does not provide direct functional insight (Walker et al., 2012; White et al., 2016). On the contrary, utilizing the VMR as a drug-screening platform identifies compounds that phenotypically improve visual function. To date, visual behavior including the VMR has not been used to screen drugs to treat RP. One reason is that due to the rods are not deemed functional until around 15 dpf (Bilotta et al., 2001; Morris and Fadool, 2005; Saade et al., 2013). However, growing evidence indicates that rods may contribute to electroretinogram and visual behavior by 5 dpf (Moyano et al., 2013; Venkatraman et al., 2020). This indicates that the scotopic behavior of the zebrafish can potentially be utilized to screen drugs to treat RP.

In this study, we utilized a scotopic VMR assay utilizing the Q344X zebrafish model to screen for drugs that can treat RP. We found that this RP model exhibited a diminished scotopic VMR behavior by 7 dpf. This response was driven by rods, as confirmed by specific rod ablation. Since it has been suggested that oxidative stress in the retina acts as one of the extrinsic factors to RP progression (Punzo et al., 2012), we leveraged this assay to screen a Redox library to determine if modulating oxidative stress could improve vision and increase rod survival. Our screen uncovered Carvedilol, a β-adrenergic receptor antagonist, enhanced the Q344X zebrafish VMR and increased rod number. We provided evidence that this drug acted on rods autonomously. Since Carvedilol is already approved by the FDA to treat heart failure and high blood pressure, this drug can easily be repurposed for the treatment of RP.

## Results

### VMR Assay Utilization for Drug Screening on Q344X Zebrafish with Scotopic Illumination

We utilized the VMR assay to screen drugs with the Q344X zebrafish model using a scotopic light stimulus. This model was selected for our drug screening as its rods begin to degenerate at 5 dpf, and the rod degeneration becomes severe by 7 dpf (Nakao et al., 2012). This rapid rod degeneration facilitates rapid evaluation of many compounds on many individual larvae. To determine the visual consequences of rod degeneration in the Q344X zebrafish, their VMR were measured under scotopic light illumination. An appropriate scotopic intensity was identified by systematically attenuating light intensity with neutral density filters until the light was 0.01 lux. To conduct the VMR assay, Q344X transgenic larvae were identified and sorted at 2 dpf by nose fluorescence. These larvae were dark adapted overnight at 6 dpf in a 96-well plate, and their VMR assessed at 7 dpf. At 7 dpf, these larvae were acclimated to the machine in darkness for 30 minutes, exposed to the scotopic light of 0.01 lux for 60 minutes, and then exposed to darkness again (Fig. 1A). The larval displacement was recorded per second for the duration of the experiment. When exposed to a light intensity of 0.01 lux, wild-type (WT) larvae displayed a robust startle response immediately after light offset (light-off VMR), while Q344X larvae displayed a significantly diminished light-off VMR (Fig. 1B). Specifically, WT larvae traveled significantly further on average than the Q344X larvae (*µ* ± standard error of the mean *(s*.*e*.*m)*: 0.281 ± 0.036 cm vs. 0.127 ± 0.031 cm) one second after light offset (Fig. 1C). Both Q344X and WT larvae did not show a response to the light onset at 0.01 lux, or they both displayed a similar response at higher intensities (Fig. S1). These results indicate that the expression of Q344X *RHO* diminished the light-off VMR of Q344X larvae at 0.01 lux.

**Figure 1:**
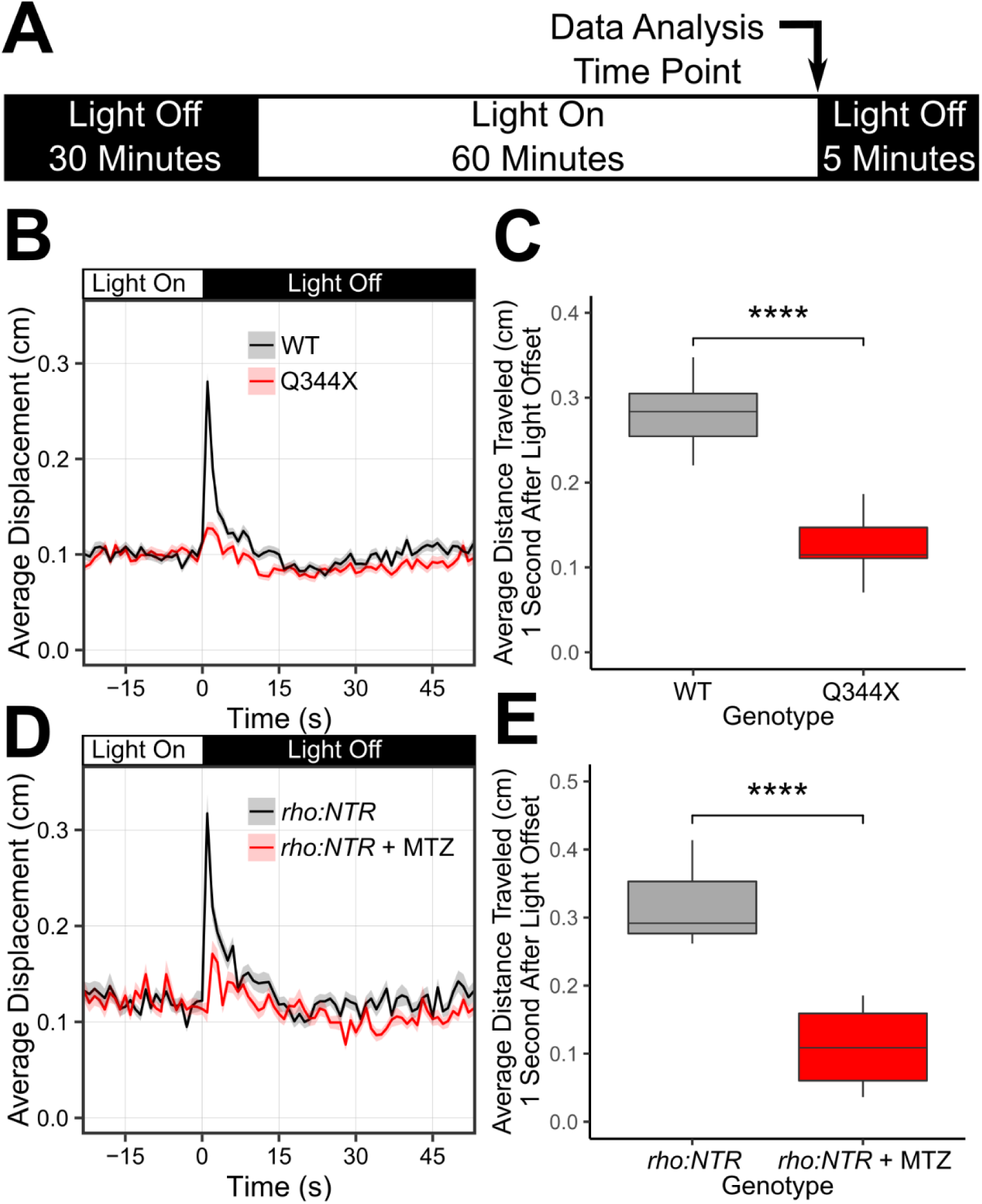
The Q344X larvae displayed diminished a scotopic light-off VMR driven by rods. (A) Schematic of the VMR protocol. On 7 dpf, larvae were habituated to the machine in the dark for 30 minutes. Then, the light stimulation was turned on and the plate was illuminated for 60 minutes. After that, the light was turned off. In this study, we mainly analyzed the VMR at light offset (light-off VMR) as indicated by the arrow. (B) The light-off VMR of wildtype (WT, black trace) and Q344X (red trace) larvae at 0.01 lux. The light was turned off at Time = 0. Each trace shows the average larval displacement of 18 biological replicates with 48 larvae per condition per replicate. The corresponding color ribbon indicates ± 1 standard error of the mean (s.e.m.). (C) Boxplot distribution of the average larval displacement of WT and Q344X larvae one second after light offset. The average displacement of WT larvae (µ ± s.e.m.): 0.281 ± 0.036cm, N = 18) was significantly larger than that of Q344X larvae (0.127 ± 0.031cm, N = 18)(Welch’s Two Sample t-test, T = 13.2, df = 33.2, p-value < 0.0001). To confirm this scotopic VMR was driven by rods, we chemically-ablated rods in larvae and subjected them to the same scotopic VMR assay. (D) The light-off VMR of larvae with nitroreductase-expressing rods treated with metronidazole (*rho:NTR* + MTZ, red trace) and without metronidazole (*rho:NTR*, black trace). Each trace shows the average displacement of 6 biological replicates with 24 larvae per condition per replicate. The corresponding color ribbon indicates ± 1 s.e.m. (E) Boxplot distribution of the average displacement of *rho:NTR* and *rho:NTR* + MTZ larvae one second after light offset. The average displacement of untreated *rho:NTR* larvae (µ ± s.e.m.): 0.317 ± 0.061cm, N = 6) was significantly larger than that of *rho:NTR* + MTZ larvae (0.110 ± 0.062cm, N = 6) (Welch’s Two Sample t-test, T = 5.9, df = 10, p-value < 0.0001).

The diminished VMR of Q344X larvae was likely caused by rod degeneration. This is plausible, as recent works detected rod ERG (Moyano et al., 2013), and rod-mediated VMR and optokinetic response (OKR) in fish larvae as early as 6 dpf (Venkatraman et al., 2020). In this study, we further confirmed rods were responsible for the diminished scotopic VMR of Q344X larvae by rod ablation. To this end, we utilized a zebrafish line expressing *nitroreductase* (*NTR*) specifically in rods under the control of the *rhodopsin* promotor (*rho:NTR*) (Walker et al., 2012). This enzyme would convert a prodrug metronidazole (MTZ) into cytotoxic substance and specifically ablate rods. In this study, the NTR-expressing larvae were treated with 2.5mM MTZ (*rho:NTR* + MTZ) from 5 dpf to 7 dpf, and their scotopic light-off VMR was measured at 7 dpf. Like the Q344X line, the rod-ablated larvae showed a significantly diminished light-off VMR compared with the untreated larvae (Fig. 1D). The average displacement of *rho:NTR* group (0.317 ± 0.061 cm) was significantly further than that of *rho:NTR* + MTZ group (0.110 ± 0.062 cm) (Fig. 1E). The reduction of scotopic light-off VMR by rod ablation indicates that the response was substantially driven by rods. This scotopic light-off VMR was then used to screen drugs that might improve rod response with the Q344X zebrafish model.

### Drug Screening Revealed that Carvedilol Ameliorates the Attenuated Q344X VMR

One of the prominent theories about RP pathogenesis is oxidative stress (Punzo et al., 2012). Since attenuating such stress might slow or prevent RP progression, we chose to screen and ENZO SCREEN-WELL® REDOX library against the Q344X zebrafish model. We chose to begin drug treatment at 5 dpf to find drugs that can ameliorate the attenuated Q344X scotopic light-off VMR because rod degeneration in this model begins at this stage. 5-dpf larvae were exposed to compounds in this library dissolved in E3 media at a final concentration of 10 μM (Wiley et al., 2017), and their scotopic light-off VMR was tested at 7 dpf. The drugs of the library come dissolved in DMSO, thus all control larvae were treated with a matching concentration of 0.1% DMSO. All larvae were maintained in the same drug solution throughout the experiment. Each drug was tested twice using embryos collected on different dates. Of the 84 drugs tested, 16 were toxic to the zebrafish at 10 μM. The VMR of the remaining 68 drug-treated larvae was normalized (Xie et al., 2019) and then ranked based on the following selection criteria: Firstly, the two biological replicates must be consistent. The consistency was determined by a High-Dimensional Nonparametric Multivariate Test (Chang et al., 2017) between the replicates. A small p-value would indicate the replicates were dissimilar, whereas a high p-value would indicate the replicates were similar. A cut off p-value of 0.9 was chosen in this study to select those replicates that were highly similar to each other. Secondly, the drug-treated VMR must be significantly different from the DMSO-treated VMR, as determined by the Hotelling’s T-squared test (Liu et al., 2015). These criteria were applied to two timeframes: just 1 second after light offset to capture immediate response, and from 1 to 30 seconds after light offset to capture changes in any of the components of the VMR (Table 1). In the 1-second timeframe, 5 drug treatments gave rise to consistent larval behavior, but none of these drug treatments gave rise to a larval VMR that was significantly different from that displayed by DMSO-treated Q344X. However, in the 30-second timeframe, four drug treatments gave rise to a consistent larval behavior, and one drug treatment, carvedilol, provided both a consistent and significant change from the DMSO-treated Q344X VMR. Carvedilol-treated Q344X exhibited a sustained scotopic light-off VMR compared with the DMSO-treated controls (Fig. 2A). To determine if carvedilol was working through the retina, eyeless *chokh/rx3* zebrafish (Loosli et al., 2003) were treated with the drug and their VMR was assessed. The *chokh/rx3* larvae did not display a light-off VMR with or without carvedilol, indicating the carvedilol acted through the retina (Fig. 2B). Previous work with the Q344X line has shown that treatment with the ADCY inhibitor SQ22536 at a concentration of 100 μM improved rod survival (Nakao et al., 2012). To determine if this rod survival can translate into improved vision, the Q344X larvae were treated with 100 μM SQ22536 from 5 dpf to 7 dpf and their scotopic light-off VMR was assessed at 7 dpf. The inhibitor was not able to increase the Q344X VMR (Fig. 2C). Our screen therefore identified carvedilol, a drug that could functionally improve the vision of the Q344X RP model.

**Table 1:**
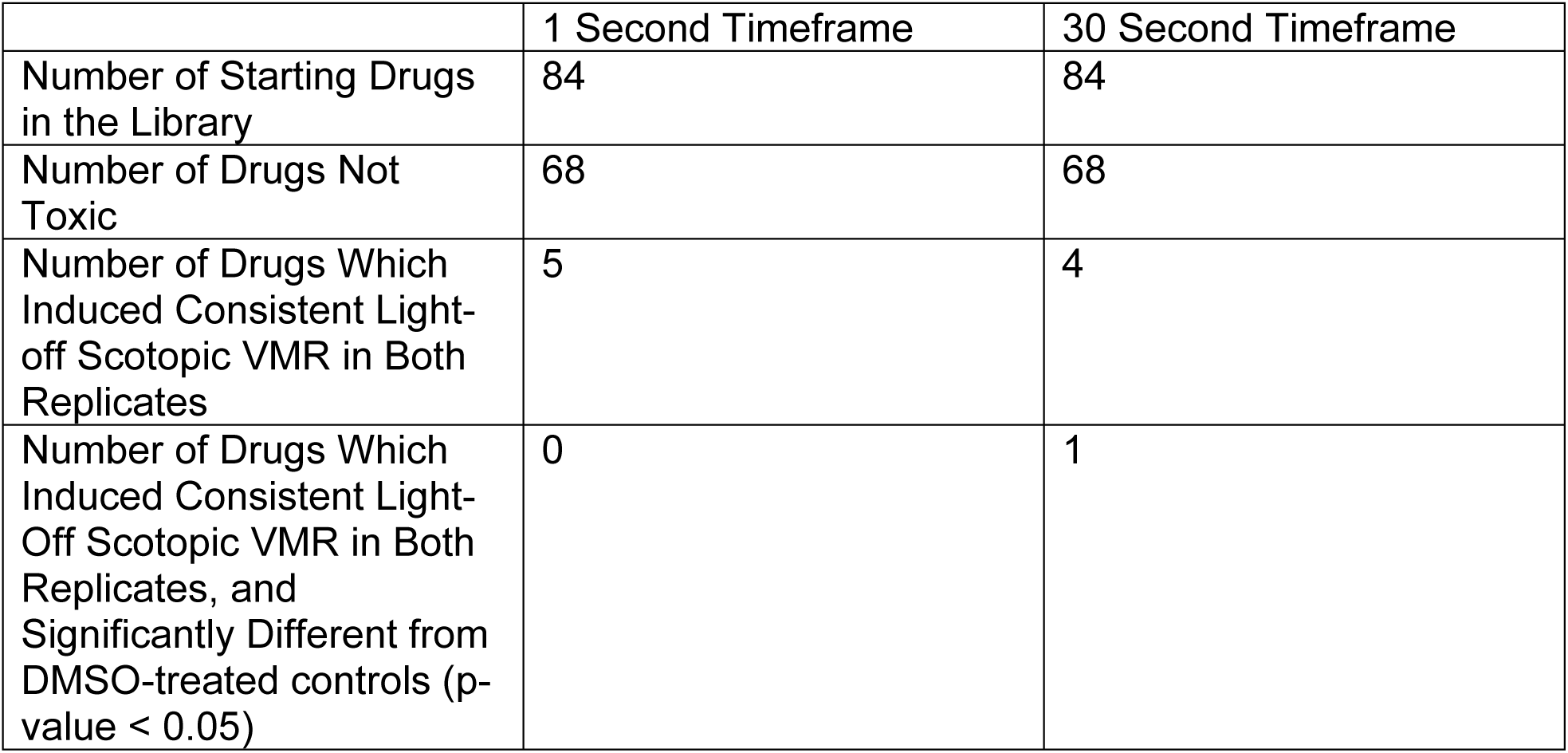
Summary of Drug-screening results. The 84 drugs in the ENZO Redox library were each applied to the Q344X larvae at 10 μM (N = 24 larvae) in two independent replicates. Of these 84 drugs, 16 were toxic. The two replicates were then compared to each other with a High-Dimensional Nonparametric Multivariate Test to determine similarity in either 1-second or 30-second timeframe. The drugs that induced consistent light-off scotopic VMR were compared to DMSO-treated Q344X controls to determine if they caused a significant change in behavior (p-value < 0.05). In the 1-second timeframe, no drugs met all criteria, but in the 30-second timeframe, one drug (carvedilol) was both consistent in both replicates and caused a significant change in light-Off scotopic VMR compared to controls.

**Figure 2:**
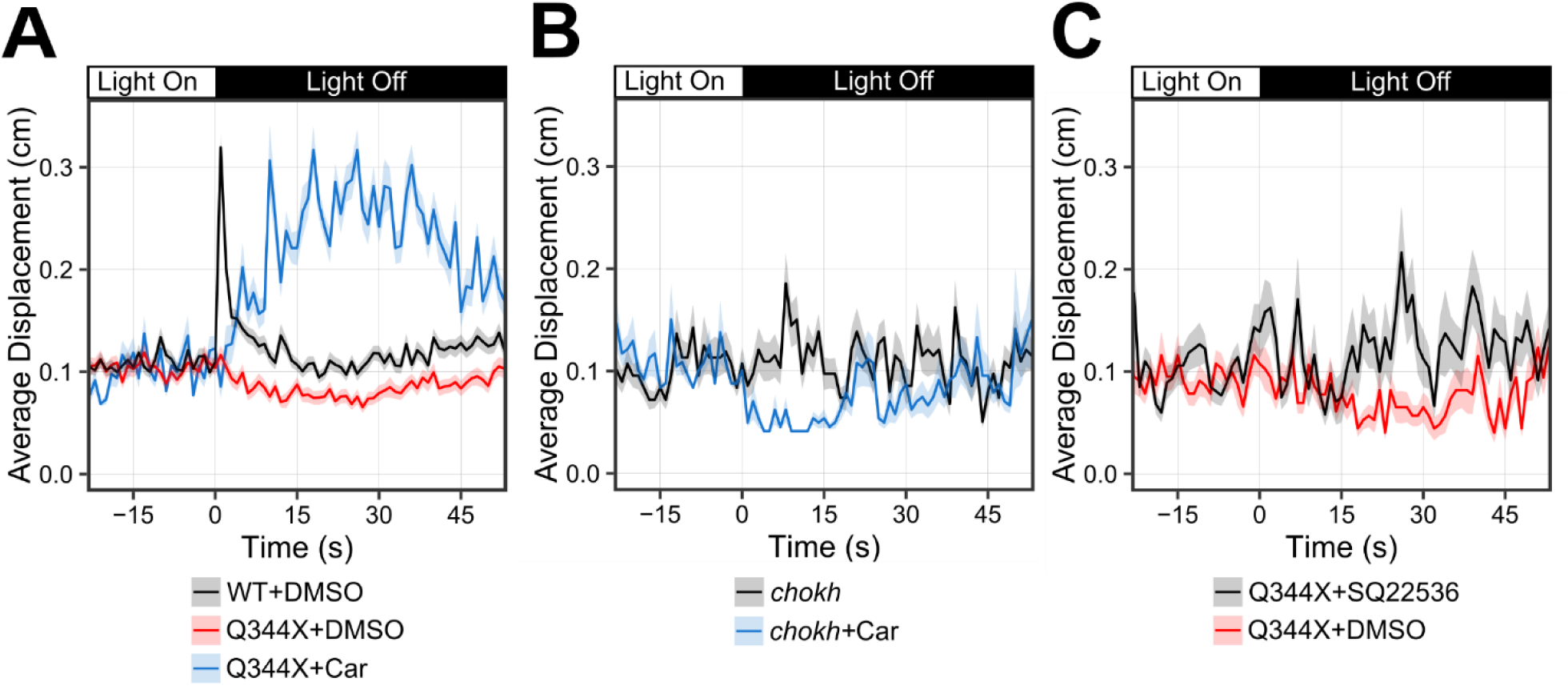
Drug screening on the Q344X zebrafish identified carvedilol as a beneficial drug. (A) Carvedilol treatment on Q344X larvae resulted in a sustained scotopic light-Off VMR (blue trace, N = 2 replicates of 24 larvae) compared to that of both DMSO-treated WT larvae and DMSO-treated Q344X larvae (black and red trace respectively, N = 9 replicates of 48 larvae). The two carvedilol replicates were highly consistent and not different from each other (High-Dimensional Nonparametric Multivariate Test, N = 24, T_HD_ = 1.78, p-value = 0.91). Each replicate demonstrated a significant change in behavior for the duration of 30 seconds after light offset above DMSO-treated Q344X larvae (Hotelling’s T-squared test, N = 24, T = 378.0 and 456.0, df = 30, p-value < 0.0001 for each replicate). (B) To determine if carvedilol’s effects are elicited through the retina, eyeless *chokh* fish were treated with carvedilol (blue trace) and their VMR was compared with untreated control (black trace). Carvedilol treatment did not increase the *chokh* VMR (Hotelling’s T-squared test, N = 24 larvae, T = 37.8, df = 30, p-value = 0.946). (C) Q344X larvae were treated with the adenylyl cyclase (ADCY) inhibitor SQ 22536 (black trace) at 100 μM to determine if inhibiting ADCY would improve the VMR compared to DMSO treatment (red trace). Treatment with SQ 22563 did not significantly improve the Q344X VMR (Hotelling’s T-squared test, N = 24 larvae, T = 40.7, df = 30, p-value = 0.952).

### Carvedilol Treatment Increased Rod Number in the Q344X Retina

Since carvedilol enhanced the scotopic VMR of the Q344X larvae and acted through the retina, it likely exhibited benefits on the degenerating rods. The drug effect on rods was evaluated by quantification of *rho:EGFP*-positive cells on wholemount and sectioned retinae (Fig. 3). On cryosections, Q344X larvae exhibited significant rod degeneration on 5 dpf at which point they were treated with carvedilol. Carvedilol-treated Q344X larvae show increased rod number in the retina compared to DMSO-treated Q344X larvae on 6 dpf and 7 dpf (Fig. 3A–D). Next, to determine the anatomical distribution of the increased number of rods in the Q344X retina, whole-mount retinae were imaged to assess rod distribution. WT larvae had a high density of rods in the lateral retina and ventral patch on 7 dpf while Q344X exhibited excessive rod degeneration in these areas (Fig. 3E). Carvedilol-treated Q344X showed an increased number of rods in both the lateral retina and the ventral patch. To quantify these observations, WT, Q344X, and carvedilol-treated Q344X were binned into three classifications based on the distribution of EGFP signal: Strong, Intermediate, and Weak (Table 2). All WT larvae were classified as Strong. The carvedilol-treated Q344X larvae had significantly more Intermediate phenotypes in the lateral and ventral views compared to the DMSO-treated Q344X group. No larvae from the carvedilol or DMSO-treated Q344X groups was classified as Strong. The correlation between rod increase and enhanced light-off VMR of Q344X larvae suggests that the extra rods mediated the visual improvement.

**Table 2:** Rod analysis on whole-mount eyes. All larvae were bleached and examined from the lateral and ventral sides. The signal were classified into 3 categories by the extent of *rho:EGFP* fluorescence. The Strong group contains the samples with high rod number/signal intensity in the lateral retina and ventral patch; the Intermediate group contains the samples with distinct rods in the lateral retina with noticeable gaps, and some rods in the ventral patch extending medially; and the Weak group contains the samples with sparse rods in the lateral retina and the most lateral edge of the ventral patch. A representative image of Strong, Intermediate and Weak type can be found in the left, middle and right column in Fig. 3E respectively. Carvedilol treatment increased the number of Q344X larvae with Intermediate phenotypes and reduced the number of Weak phenotypes in both the lateral (Chi-square test, *χ*^2^ = 4.09, df = 1, p-value < 0.05) and ventral views (Chi-square test, *χ*^2^ = 5.33, df = 1, p-value < 0.05). No Q344X larvae was classified as Strong with or without carvedilol treatment.

|  | Strong | Intermediate | Weak |
| --- | --- | --- | --- |
| WT Lateral | 10 | 0 | 0 |
| Q344X Lateral | 0 | 9 | 15 |
| Q344X+Car Lateral | 0 | 16 | 8 |
| WT Ventral | 10 | 0 | 0 |
| Q344X Ventral | 0 | 8 | 16 |
| Q344X+Car Ventral | 0 | 16 | 8 |

**Figure 3:**
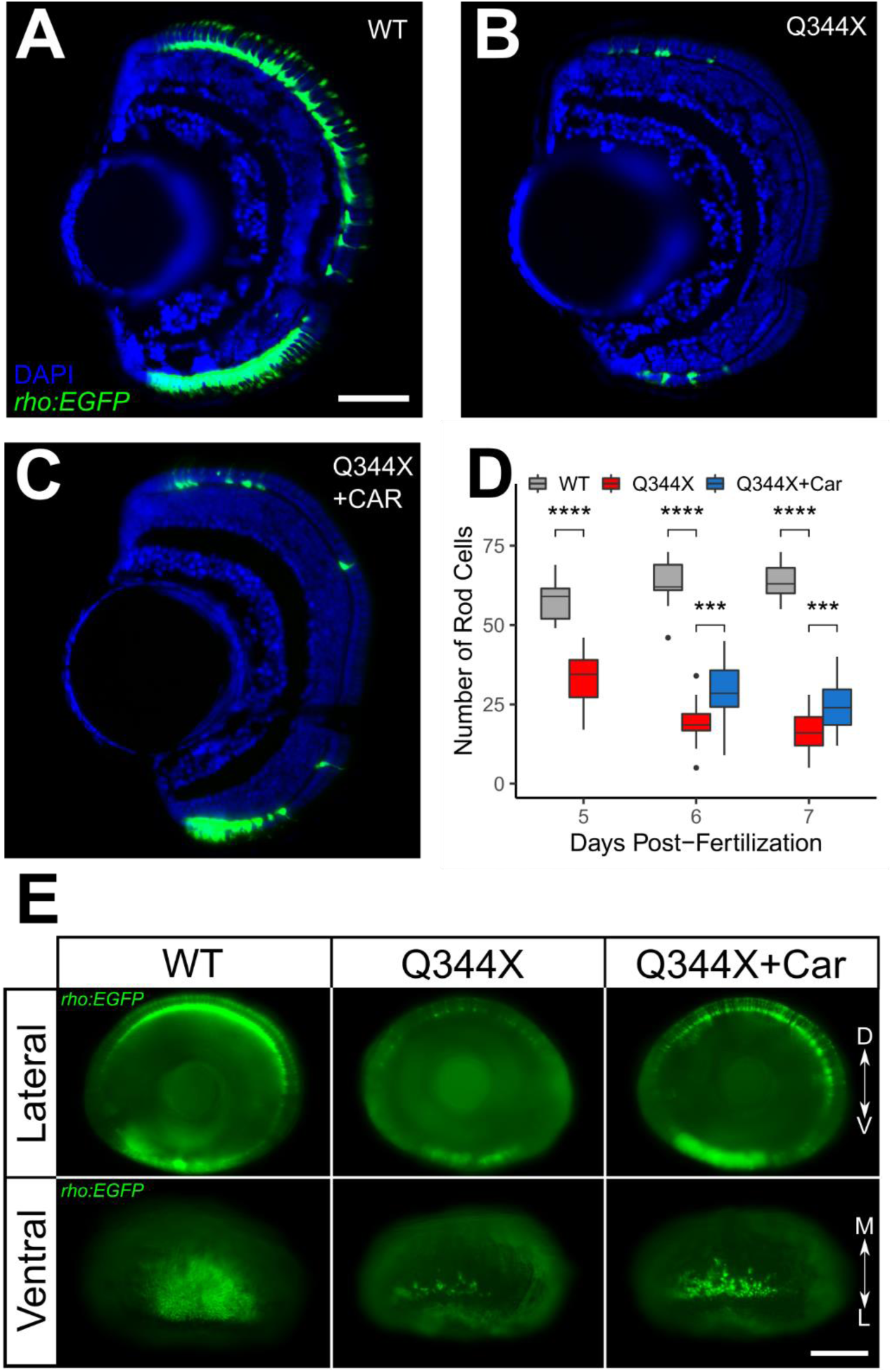
Carvedilol treatment increased rod numbers in the Q344X larvae. Representative retinal cryosection of a (A) wildtype (WT), (B) Q344X, and (C) carvedilol-treated Q344X (Car) larva at 7 dpf. Rods were labeled by EGFP expression driven by *rho* promoter (*rho:EGFP)*, and the nuclei were counterstained with DAPI. Scale = 50 μm. (D) Quantification of rod number in WT, Q344X, and carvedilol-treated Q344X retinal cryosections from 5 dpf to 7 dpf. There was a statistically significant difference in rod number between groups at all stages determined by one-way ANOVA at 5 dpf (WT, N = 11; Q344X, N =16; *F*(1,25) = 71.04, p-value < 0.0001), at 6 dpf (WT, N = 9; Q344X, N =20; Q344X+Car, N =21 ; *F*(2,44) = 96.9, p-value < 0.0001), and at 7 dpf (WT, N = 9; Q344X, N =17; Q344X+Car, N =11; *F*(2,41) = 167.9, p-value < 0.0001). The effect of Q344X rod degeneration and carvedilol treatment on rod number was assessed post hoc by pairwise t-test with false discovery rate correction at 6 dpf (WT – Q344X, p-value < 0.0001; Q344X – Q344X+Car, p-value < 0.001) and at 7 dpf (WT – Q344X, p-value < 0.0001; Q344X – Q344X+Car, p-value < 0.001). E) Representative whole-eye images of WT, Q344X, and carvedilol-treated Q344X larvae at 7 dpf. Rods were labeled by EGFP expression. Left column: WT rods were mainly found on dorsal and ventral retina (top). They were abundantly present in the ventral patch of the retina extending medially (bottom). Middle column: Q344X rods were mostly degenerated at the same stage (top). There were only a handful of rods remaining near the lateral edge of the Q344X retina (bottom). Right column: Carvedilol treatment increased the number of Q344X rods on both dorsal and ventral retina (top); however, gaps of missing rods were still apparently on dorsal retina. More rods were observed in the ventral patch of the carvedilol-treated retina (bottom). Scale = 100 μm. D = Dorsal, V = Ventral, M = Medial, L = Lateral.

The extent of Q344X rod number improvement was evaluated with a longer carvedilol treatment period. Q344X larvae were treated with 10 μM carvedilol beginning at 3 dpf. Given the earlier developmental stage of the larvae, the drug and media were refreshed daily to maintain the health of the larvae. Treatment at 3 dpf was compared to treatment at 5 dpf to determine if earlier carvedilol treatment is more effective. There was no difference in rod number between any of the Q344X and WT groups at 3 dpf and 4 dpf indicating that Q344X rod degeneration is not significant at these stages (Fig. 4). Q344X rod degeneration does become significant at 5 dpf, and the earlier carvedilol treatment beginning at 3 dpf significantly increased the rod number at 5 dpf (Fig. 4). Carvedilol treatment beginning at 5 dpf with daily refreshment still improved rod number in the Q344X zebrafish at 6 dpf and 7 dpf, however carvedilol treatment beginning at 3 dpf resulted in significantly higher rod numbers than the later 5 dpf treatment (Fig. 4.). These results suggest that earlier carvedilol drug treatment improves the number of Q344X rods better than later treatment.

**Figure 4:**
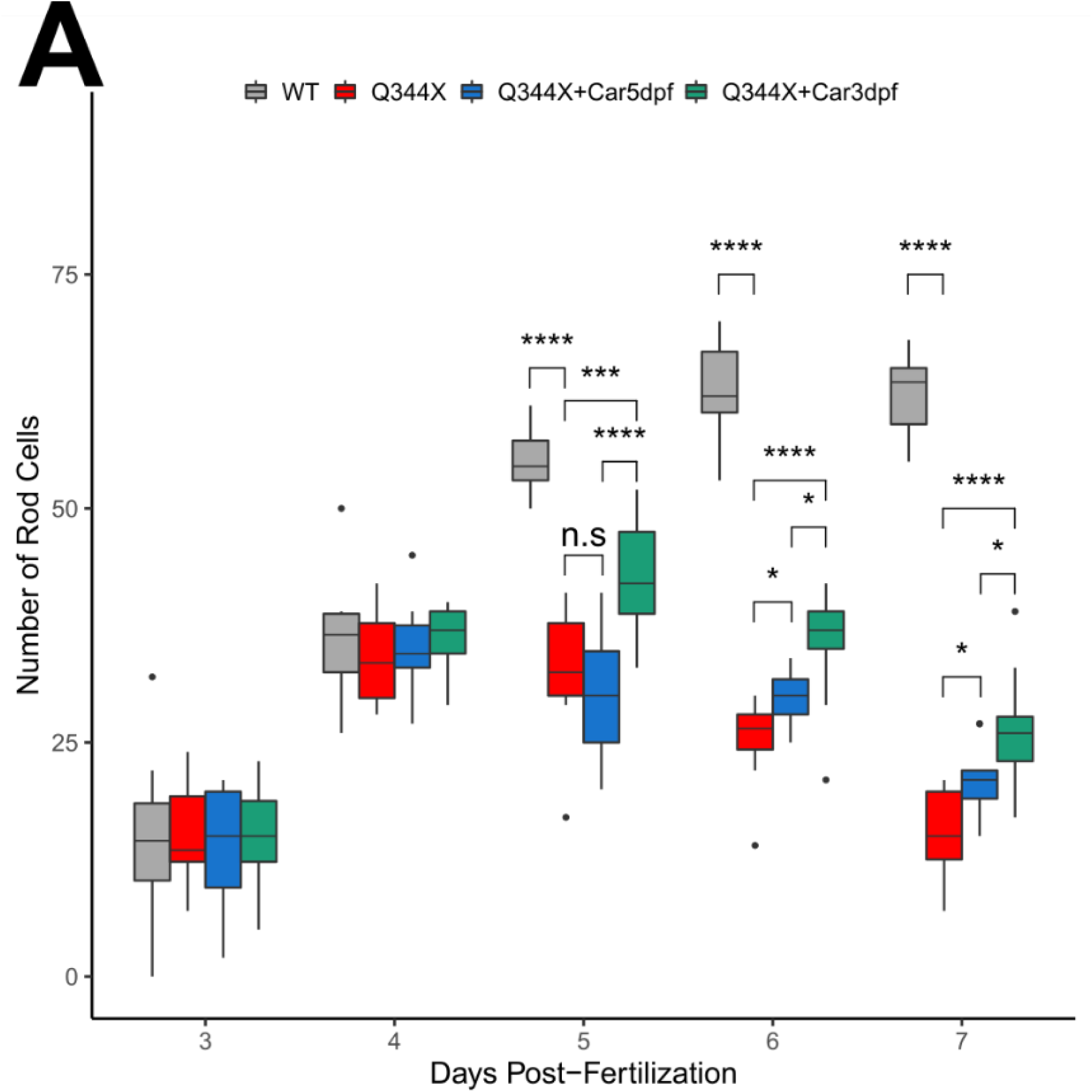
Carvedilol treatment beginning at 3 dpf increased rod numbers in the Q344X larvae greater than treatment beginning at 5 dpf. (A) Quantification of rod number in WT, Q344X, and Q344X treated with carvedilol beginning at 3 dpf and 5 dpf. retinal cryosections from 3 dpf to 7 dpf. There was no statistically significant difference in rod number between groups at stages 3 dpf and 4 dpf determined by one-way ANOVA (3 dpf; N = 10; *F*(3,36) = 0.1, p-value = 0.95); (4 dpf; N = 10 ; *F*(3,36) = 0.5, p-value = 0.69). There was a statistically significant difference in rod number between groups at stages 5 dpf through 7 dpf determined by one-way ANOVA at 5 dpf (N = 10; *F*(3,36) = 0.1, p-value < 0.0001), at 6 dpf (N = 10; *F*(3,36) = 0.1, p-value < 0.0001) and at 7 dpf (N = 10; *F*(3,36) = 0.1, p-value < 0.0001). The effect of Q344X rod degeneration and carvedilol treatment on rod number was assessed post hoc by pairwise t-test with false discovery rate correction at 5 dpf (WT – Q344X, p-value < 0.0001; Q344X – Q344X+Car3dpf, p-value < 0.001; Q344X – Q344X+Car5dpf, p-value = 0.36, Q344X+Car3dpf – Q344X+Car5dpf, p-value < 0.0001), at 6 dpf (WT – Q344X, p-value < 0.0001; Q344X – Q344X+Car3dpf, p-value < 0.0001; Q344X – Q344X+Car5dpf, p-value < 0.05; Q344X+Car3dpf – Q344X+Car5dpf, p-value < 0.05), and at 7 dpf (WT – Q344X, p-value < 0.0001; Q344X – Q344X+Car3dpf, p-value < 0.0001; Q344X – Q344X+Car5dpf, p-value < 0.05; Q344X+Car3dpf – Q344X+Car5dpf, p-value < 0.05).

### Carvedilol can Inhibit β-adrenergic Signaling in Rod-like Y79 Retinoblastoma Cells

Carvedilol is a β-blocker that that binds to β-adrenergic receptors and inhibits adrenergic signaling. However, its retinal target is unknown, and it may act directly on rods. To evaluate this possibility, we examined the effect of carvedilol treatment on a rod-like cell line, the Y79 human retinoblastoma line which uniquely expresses rod-specific genes (Di Polo and Farber, 1995). The level of adrenergic signaling was determined by GPCR-modulated changes in cAMP levels as measured by a cAMP-sensitive luciferase. First, the Y79 cells were transfected with the luciferase reporter, and then they were exposed to half-log dilutions of isoproterenol, a β-adrenergic receptor agonist. Isoproterenol was capable of inducing cAMP signaling in the transfected Y79 cells with a pEC50 of 7.5 ± 1.1 (Fig. 5A). The cAMP level was not increased in controls treated with matching DMSO percentage to dissolve isoproterenol. The relative cAMP level did not increase much above 10 μM isoproterenol. To determine if carvedilol treatment can inhibit this isoproterenol-mediated cAMP increase, the transfected Y79 cells were pretreated with half-log dilutions of carvedilol and then challenged with a dose of 10 μM isoproterenol that would induce saturating relative cAMP level according to Fig. 5A. Carvedilol pretreatment was able to prevent isoproterenol-mediated cAMP signaling with a pIC50 of 6.5 ± 0.7 (Fig. 5B). Therefore, carvedilol likely bound to the β-adrenergic receptors and directly elicited its beneficial effects on rods.

**Figure 5:**
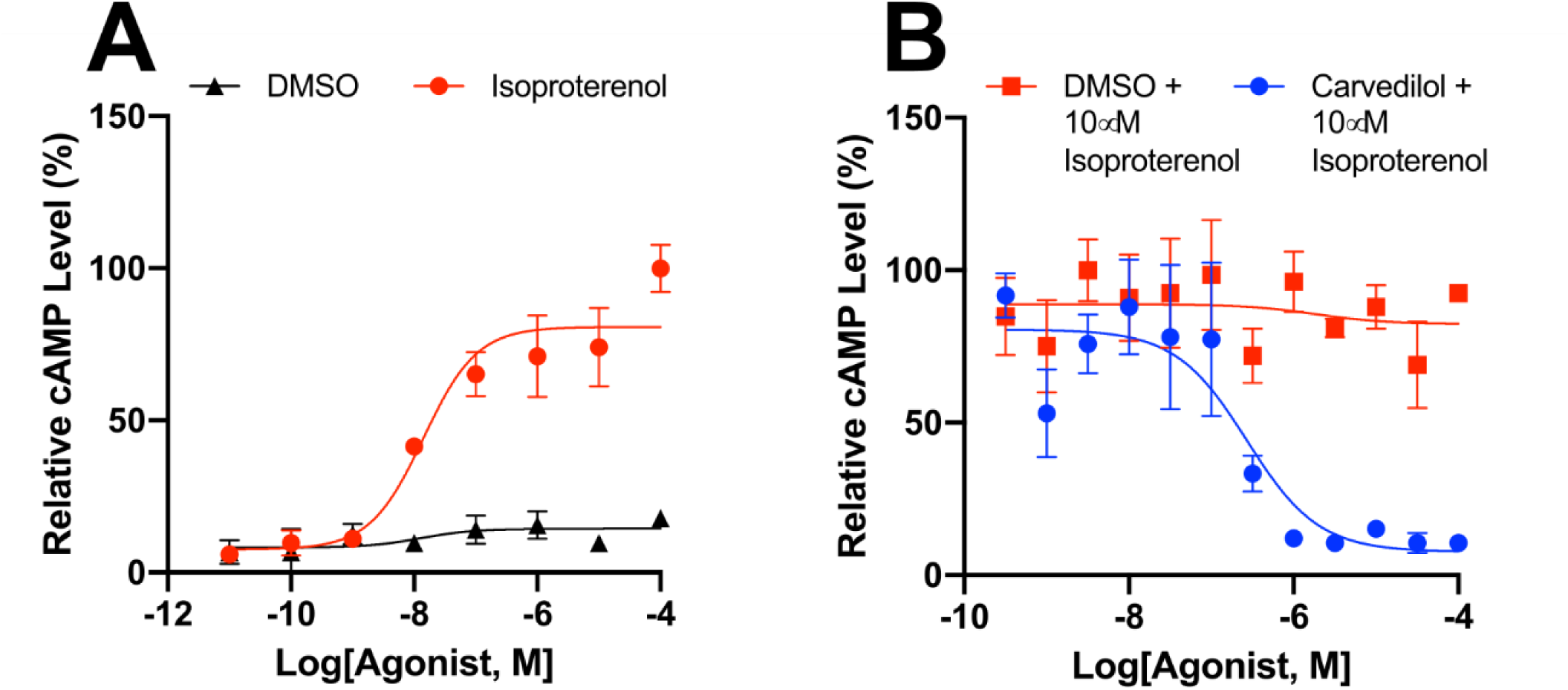
Carvedilol treatment might directly act on rods cells. To determine the extent to which carvedilol act directly on rods, we conducted a GloSensor cAMP assay with rod-like human Y79 cells. (A) Representative dose-response curves of GloSensor-transfected Y79 cells treated with half-log concentrations of isoproterenol (red trace; N = 4) or percentage-matched DMSO (black trace; N = 4). These plots were normalized to the maximum average luminescent level recorded per experiment. Error bars show ± 1 s.e.m. Isoproterenol was capable of increasing cAMP signaling through β-adrenergic receptor binding with an pEC50 of 7.49 ± 1.07. (B) Representative dose-response curves of GloSensor-transfected Y79 cells pretreated with half log doses of carvedilol (blue trace; N = 4) or percentage-matched DMSO (red trace; N = 4). Cells were then challenged with a 10 μM isoproterenol that could induce maximal cAMP response, as shown in Fig. 5A. Carvedilol pretreatment was capable of preventing isoproterenol-mediated cAMP increases with an pIC50 of 6.51 ± 0.67.

## Discussion

There are no approved cures to treat RP. To address this unmet need, we have utilized the Q344X RP zebrafish model to find drugs that can help patients suffering from the disease. We found that rod degeneration in the Q344X zebrafish resulted in a deficient scotopic light-off VMR, a locomotor response displayed during drastic light offset. This behavior was driven by rods, as it was diminished by chemical ablation of the rods. We leveraged this behavior as a functional assay to screen for beneficial drugs that could enhance the response in the Q344X model. We found that both the scotopic VMR and retinal histology of the Q344X model were improved by carvedilol. Since this drug is approved by the FDA, it can potentially expedite the development of a new treatment for RP patients.

Carvedilol has several known modes of action. It is primarily classified as a β-blocker; however, it has also been demonstrated to act as an α1-blocker, a calcium channel agonist at high concentration, and a free radical scavenger (Giannattasio et al., 1992; McTavish et al., 1993). Carvedilol may mediate its visual benefit through some of these pathways. Traditionally, β-blockers are seen only as antagonists that prevent epinephrine from binding β-adrenergic receptors. Epinephrine is present in the mouse subretinal space and increases with light exposure (Hadjiconstantinou et al., 1983). Blocking epinephrine signaling can potentially lower cAMP levels in the Q344X rods by preventing endogenous ADCY signaling. Interestingly, carvedilol also acts as an atypical β-blocker which is capable of inducing biased signaling (Wisler et al., 2007). Specifically, carvedilol can promote β-arrestin signaling while acting as an inverse agonist towards G protein signaling (Wang et al., 2017). This type of β-arrestin signaling has been shown to have anti-apoptotic effects (Ahn et al., 2009; DeFea et al., 2000; Povsic et al., 2003) that may prevent Q344X rod death. Carvedilol may also exert protective effects on Q344X rods through α1 blockade since selectively blocking Gq-coupled α1 adrenergic receptors can prevent photoreceptor degeneration in a Stargardt Disease model (Chen et al., 2013; Chen et al., 2016). In our study, it is unlikely that carvedilol is acting as a calcium channel blocker at the tested dose since it would likely stop the larval heart beating and kill the larva before its VMR could be measured (Chopra et al., 2010; Stainier, 2001). Since our study did not identify any other beneficial compounds from this REDOX library, carvedilol probably did not exert its visual benefit on the Q344X model as a radical-scavenging antioxidant.

More evidence is available which suggests that β-blockers may be able to treat RP. A recent study has identified that another β-blocker, metipranolol, is capable improving rod survival and electroretinogram in the *rd10* mouse (Kanan et al., 2019). Another study has found that the β-blocker metoprolol can provide protection against bright light-induced retinal degeneration, and metoprolol protection can be increased by co-treatment with other GPCR agonists and antagonists (Chen et al., 2016). Carvedilol has already been shown to have beneficial effects with treating other eye-disease models. Carvedilol can lower intraocular pressure (IOP) in the eye of rabbits (Nakaya et al., 2012). Also, carvedilol has neuroprotective effects on retinal ganglion cells in an optic nerve injury mouse model (Liu and Liu, 2019). However, it is unknown if β-blockers such as carvedilol can work directly on rods, so we employed the human rod-like Y79 line to determine this. We were able to demonstrate that carvedilol can inhibit isoproterenol-mediated G protein signaling in the Y79 cells. In addition, one of carvedilol’s target receptors, the β-_1_ adrenergic receptor, is expressed in mouse rods (Siegert et al., 2012). These results suggest that carvedilol is able to elicit its therapeutic effects directly on the rods, and it targets adrenergic signaling in the eye that may treat RP.

Targeting GPCR signaling through adrenergic receptors is an attractive method for the treating Q344X RP. While the full disease mechanism is unknown, it is believed that mislocalized activation of rhodopsin in the inner segment of Q344X rods induces ADCY activation resulting in cAMP increase and apoptosis (Concepcion and Chen, 2010; Nakao et al., 2012; Portera-Cailliau et al., 1994; Sung et al., 1994). This highlights ADCY as a potential drug target. Previous work with the Q344X zebrafish, as well as the Stargardt Disease mouse, has shown that inhibition of ADCY with the inhibitor SQ 22536 improved photoreceptor survival (Chen et al., 2013; Nakao et al., 2012). However, SQ 22536 treatment did not improve the VMR displayed by Q344X larvae. Therefore, the improved Q344X VMR from carvedilol treatment may be due to other chemical properties of the drug.

We have performed the first functional drug screen for RP which has discovered carvedilol as a positive hit. The common starting concentration of 10 μM was chosen to minimize toxic effects while maximizing the chances of finding an effective dose (Wiley et al., 2017). Drug screening with the Q344X zebrafish model begun at 5 dpf due to the onset of rod degeneration and the display of a variety of visual behaviors including the VMR and OKR (Chhetri et al., 2014; Easter and Nicola, 1997; Emran et al., 2007; Liu et al., 2015). This stage also was chosen to allow the larvae to developmentally mature to a stage as far as possible to minimize potential toxicity from drugs. Drugs identified through this screening method may provide a beneficial result, but it may not be the optimal treatment period. In the case of carvedilol, we tested a treatment period beginning at 3 dpf and determined that earlier treatment further improved rod survival. Utilizing the behavioral drug screen at 5 dpf and further investigating hits with earlier treatment is an efficient method for identifying the best hits for further translation. Investigating the VMR and rod survival at stages later than 7 dpf becomes more complicated because larval zebrafish deplete their yolk at around 9 dpf and require feeding to survive. Larval feeding and foraging introduce extra variability in the behavioral characterizations of drug effects. Future research will elucidate the mechanism through which these carvedilol-regulated pathways increase rod numbers and should test carvedilol’s efficacy on the Q344X mouse model (Sandoval et al., 2014) to validate the translational value of carvedilol for RP treatment. Our screen also lays the foundation for drug screening with fish modeling different classes of RP mutations.

In this study, we have demonstrated that carvedilol can increase rod number and the scotopic VMR of the Q344X RP zebrafish. Carvedilol is a unique β-blocker that can induce biased β-adrenergic signaling as well as antagonisms of α_1_-adrenergic receptors. It has already shown promise in other animal models of eye disease, and since the drug already has FDA approval, it can potentially be quickly repurposed to treat RP. Therefore, our data may provide a new drug for treating RP patients.

## Materials and Methods

### Animals

Zebrafish of the AB background were utilized for all experiments <https://zfin.org/ZDB-GENO-960809-7>. Adult and larval zebrafish were maintained and bred using standard procedure (Westerfield, 2007). Adult fish were placed in breeding tanks the night before breeding after receiving all meals. Adult fish began spawning at 9:00am, and embryos were collected before 10:30am. Larval zebrafish were reared until 7 days post-fertilization (dpf) in E3 medium in an incubator at 28°C. The fish incubator was kept on a 14hr light and 10hr dark cycle. E3 medium was changed daily, and healthy embryos were kept for experiments. All protocols were approved by the Purdue University Institutional Animal Care and Use Committee.

### Transgenic Animals

*Tg(rho:Hsa*.*RH1_Q344X)* transgenic animals were generated previously (Nakao et al., 2012) and are referred to in this study as Q344X. Q344X larvae were identified on 2 dpf through the expression of EGFP under the control of 1.1kbp promoter of *olfactory marker protein* (*omp*) contained in the transgenic cassette. Their genotype was verified via PCR with the following primers: 5′-CCAGCGTGGCATTCTACATC-3′ and 5′-AACGCTTACAATTTACGCCT-3′. The rods in the Q344X line were labeled with the *Tg(−3*.*7rho:EGFP)* transgene (Hamaoka et al., 2002) and are referred to in this study as *rho:EGFP*. Zebrafish expressing *nitrodreductase* in rod photoreceptors, *Tg(rho:YFP-Eco*.*NfsB)*^*gmc500*^, were generated previously (Walker et al., 2012) and referred to in this study as *rho:NTR*.

### Drug Treatment

The ENZO SCREEN-WELL® REDOX library was used for drug screening (ENZO Life Sciences, BML-2835-0100). Carvedilol was ordered from ENZO Life Sciences (BML-AR112-0100) for further experiments and from MilliporeSigma (C3993-50MG) for confirming the positive effects observed in specific behavioral experiments (data not shown). Unless otherwise stated, all drugs were dissolved in DMSO. The maximum DMSO percentage used was 0.1%. Thirty larvae were exposed per drug dissolved in 15 mL E3 media in a 15 mm petri dish. The treatment began on 3 dpf or 5 dpf as stated. The media were not refreshed during experiment unless otherwise stated. The treated larvae were directly transferred into the 96-well plate with their corresponding E3 medium with drugs to ensure consistent drug dosing throughout the treatment period.

### Rod Ablation in rho:NTR Zebrafish

To chemically ablate rods, we used the zebrafish line, *Tg(rho:YFP-Eco*.*NfsB)*^*gmc500*^, expressing *nitroreductase* (NTR) under the control of the rhodopsin promotor (Walker et al., 2012) (*rho:NTR)*. Treatment with the prodrug metronidazole (MTZ) specifically ablates rod photoreceptors. Specifically, NTR-expressing larvae were treated with 2.5 mM MTZ from 5 dpf to 7 dpf. Their VMR was compared with the untreated larvae on 7 dpf.

### Retinal Histology and Imaging

All larvae were fixed in 4% paraformaldehyde (PFA) overnight at 4°C. For retinal cryosections, fixed larvae were infiltrated with 30% sucrose overnight at 4°C prior to imbedding in Tissue Freezing Medium (GeneralData, TFM). Ten micrometers-thick cryosections were collected on Fisherbrand Superfrost Plus Microscope Slides (Thermo Fisher Scientific, 12-550-15). The sections containing the optic nerve were analyzed for anatomical reference.

### Whole-animal Preparation

To visualize rod distribution in the retina, PFA-fixed larvae were bleached with 1% KOH/3% H_2_O_2_ for 40 minutes at room temperature to bleach the black pigment from the retinal pigment epithelium. The bleached embryos were imbedded a 3% methyl cellulose solution for observation.

### Microscope and Camera

All samples were imaged with an Olympus BX51 microscope (Olympus) and a SPOT RT3 Color Slider camera (SPOT Imaging).

### Visual Motor Response Assay

A ZebraBox system from ViewPoint Life Sciences was utilized for the Visual Motor Response (VMR) assay. Individual zebrafish larvae were placed in 96-well plate format using Whatman UNIPlate square 96-well plates (VWR, 13503-152). In order to produce a scotopic stimulus, the ZebraBox was modified to attenuate the light intensity beyond its lowest limit by fitting neutral-density filters in the light path. Seven neutral density filters (BarnDoor Film and Video Lighting, E209R), each allowing approximately 40% transmittance, were stacked between the light source and the plate holder until a final intensity of 0.01 lux was attained. In our scotopic experiments, the machine was also powered at 5% in order to prevent instability from the LED light source. The larval displacement was collected by the tracking mode which binned the activity every second.

To conduct the scotopic VMR assay, larvae were sorted and grown in 100 x 15 mm petri dishes (VWR, 25384-088) with 15 mL E3 media in a density of 30 larvae from 2 dpf to 5 dpf. Larvae were transferred to 96-well plates with one larva per well on the morning of 6 dpf and dark adapted overnight. On 7 dpf, the dark-adapted larvae were placed in the ZebraBox and their scotopic VMR was measured. For drug screening with larvae, this procedure was the same except larvae were exposed to drugs in petri dishes on 5 dpf. In this study, the following protocol was used: 30 minutes in the dark followed by a 60-minute scotopic light illumination at 0.01 lux, and then a light offset for 5 minutes (Fig.1A). All VMR experiments were conducted between the 9am and 6pm to minimize the effect of circadian rhythm on vision (Emran et al., 2010).

### Light Stimulus Intensity

Light intensity of the ZebraBox LED spectrum was measured with a SpectriLight ILT950 Spectroradiometer (International Light Technologies). The total irradiance of the LED stimulus over the entire visible spectrum at 5% power output was 3.2 µW cm^-2^ (0.0063 µW cm^-2^ at 500 nm wavelength). The corresponding illuminance was 8.0 lux. The light intensity was further reduced by fitting neutral-density filters in the light path as described above. These neutral-density filters did not alter the color spectrum of the LED light. The light intensity with neutral-density filters was calculated by multiplying the light intensity emitted by the machine with the transmittance of each neutral-density filter. The irradiance of the final scotopic stimulus used in this study was 0.005 µW cm^-2^ (1.80e-5 µW cm^-2^ at 500 nm). The corresponding illuminance was 0.01 lux.

### Y79 Cell Culture and cAMP Assays

The human Y79 cell line was obtained from American Type Culture Collection (ATCC HTB-18). These cells were cultured in RPMI-1640 Medium (ATCC, 30-2001) with 15% Fetal Bovine Serum (ATCC, 30-2020) at 37°C with 5% CO_2_. To measure cAMP levels in the cells, the GloSensor Technology -22F cAMP plasmid (Promega, E2301) was used with GloSensor Assay Reagent (Promega, E1290). Four million cells were seeded into 10 cm dishes with 10 mL of Opti-MEM Reduced Serum Medium (ThermoFisher, 31985062) for transfection. The cells were transfected with 20µg of GloSensor plasmid utilizing X-tremeGENE HP DNA Transfection Reagent (MilliporeSigma, 6366244001) at a 2:1 ratio of plasmid to X-tremeGENE reagent, according to manufacturer’s instructions. These cells were transfected for 24 hours, and then they were transferred back into RPMI for another 24 hours. Twenty-five thousand cells were then seeded into a low-volume white 384-well plate per well (Greiner Bio-one, 784080). The cAMP assay was carried out according to the GloSensor protocol for suspension cells. Cells were either treated with DMSO or carvedilol for 20 minutes at room temperature prior to treatment with isoproterenol. Luminosity was recorded 20 minutes after drug or DMSO vehicle addition with a FlexStation 3 Multi-Mode Microplate Reader (Molecular Devices).

### Data Visualization and Statistical Analysis

#### General Data and Statistics

All standard statistical analyses were performed with R version 3.6.0 http://www.r-project.org.

#### VMR Data

Statistical analyses and data processing were performed with R version 3.6.0 http://www.r-project.org. Raw data from the VMR assay was processed and extracted by Data Workshop (ViewPoint Life Sciences). Data figures were created using *ggplot2* package in R (Wickham, 2016).

The VMR data were normalized for baseline activity, light-intensity variation per well, and batch effect (i.e. biological replicate) by linear-regression models as previously described (Xie et al., 2019). Additionally, offset values were added to the normalized activity to prevent negative displacement.

To determine if each VMR replicate from drug-treated Q344X larvae during drug screening was consistent with the other replicate, a High-Dimensional Nonparametric Multivariate Test (Chang et al., 2017). This test was chosen because the number of observations (i.e. sample size) for each VMR is less than the dimension of the dataset. The dimension is the length of the time period used in the analysis and the sample size is the number of drug-treated larvae. The High-Dimensional Hypothesis test was implemented in the R package *HDtest*.

The Hotelling’s T-squared test (Hotelling, 1931) was used to test significant changes in zebrafish displacement from 1 to 30 seconds after the light change. This test is the multivariate version of the T-test which follows the F-distribution. The test statistic for the Hotelling’s T-squared test is calculated as : 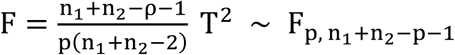 where *n*_1_ and *n*_2_ are the sample size. ρ is the dimension which is the time interval used in the analysis. The Hotelling’s T-squared test was used for VMR analysis due to a number of advantages: 1. The Type I error rate is controlled. 2. The relationship between multiple variables is considered. 3. It can generate an overall conclusion even if multiple (single) t-tests are inconsistent. The null hypothesis for the experiment is the group means for all response variables are equal which means the mean vector of the distance travelled for the two chosen groups are the same (*μ*_1_ = *μ*_2_). The Hotelling T-squared test analysis was performed on the R package *Hotelling* (Curran, 2018) with some reshape of the dataset.

#### Y79 cAMP Data

Luminosity data obtained from the Y79 cell line was analyzed and plotted using Graphpad Prism (version 8, GraphPad Software). Data were plotted with the non-linear fit method under “log(agonist) vs. response – Variable slope (three parameters)”. pEC50 (negative log of half maximal effective concentration) and pIC50 (negative log of half maximal inhibitory concentration) were calculated through the above-mentioned non-linear fit.

## Data availability

The raw zebrafish behavioral data is available on the Harvard Dataverse <https://doi.org/10.7910/DVN/JYLWH1>. The R scripts to reproduce the analyses and plots reported in this paper are available in GitHub <https://github.com/zhanzmr/Zebrafish_Model>.

## Contributions

Y.F.L., C.P.P., M. Z. (Mingzhi Zhang) and M. T. conceptualized this study. Y.F.L. and L.G. designed the experiments. M.T. generated and provided the *Tg(rho:Hsa*.*RH1_Q344X)* zebrafish line. L.G. performed drug screening with VMR. L.G. and R.J. performed rod imaging through cryosections and whole mount. J.M. and L.Z. contributed the *Tg(−3*.*7rho:YFP-nfsB)*^*gmc500*^ zebrafish line and performed metronidazole treatment. M.Z. (Mengrui Zhang), R.X., Y.C., and W.Z. analyzed the VMR data. L.G., M.J.K. and R.M.vR. performed cAMP assays on the Y79 cell line and interpreted the data. L.G., M.T. and Y.F.L. interpreted the results. L.G., and Y.F.L. drafted the manuscript. All authors contributed to the critical revision of the manuscript and approved the final version.

## Table and Figure Legends

**Supplementary Figure 1:**
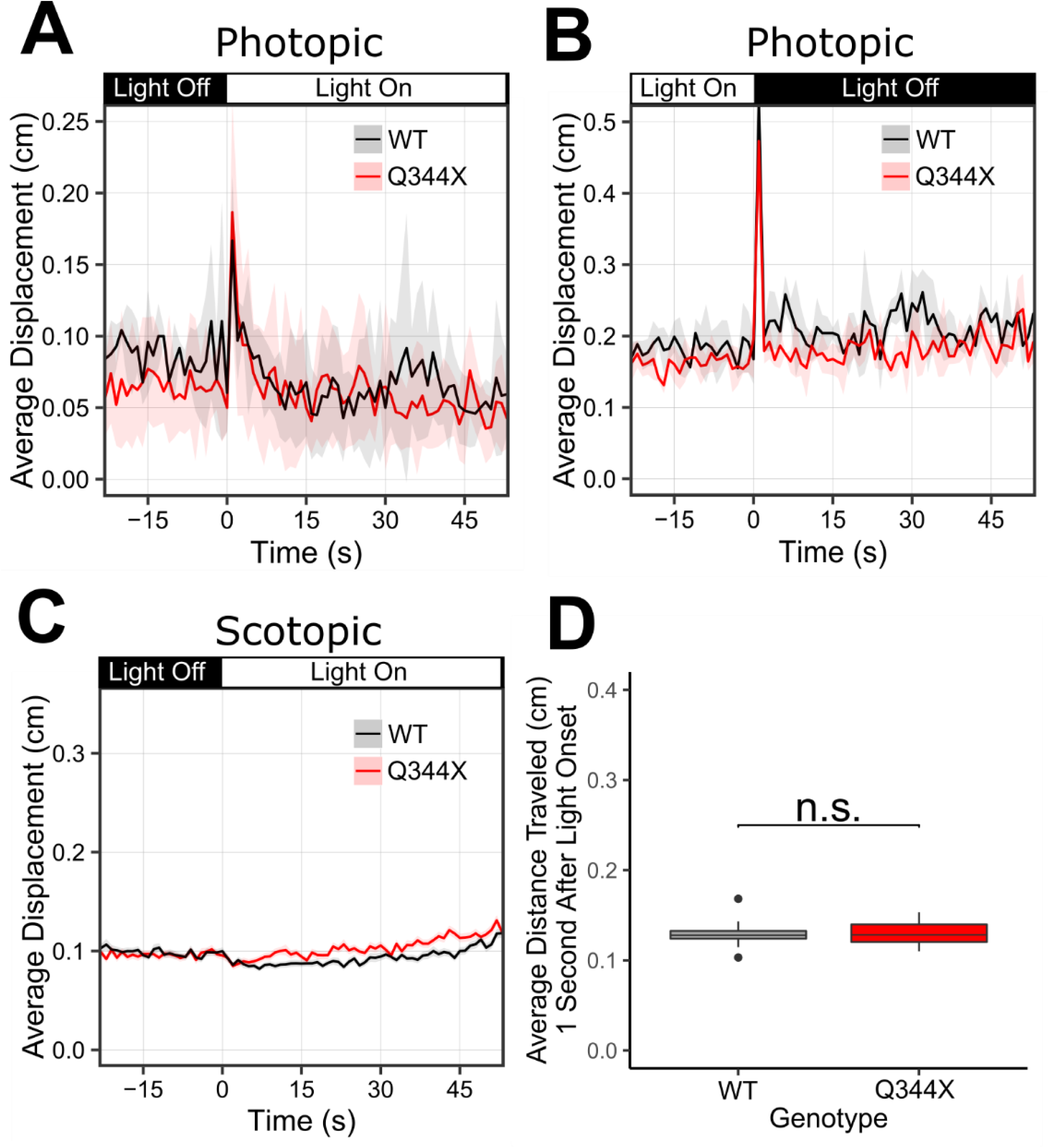
(A) Both Q344X and WT larvae respond a photopic light onset stimulus. The light-on VMR of wildtype (WT, black trace) and Q344X (red trace) larvae at 880 lux. The light was turned on at Time = 0. Each trace shows the average larval displacement of 3 technical replicates with 32 larvae per replicate. The corresponding color ribbon indicates ± 1 s.e.m. (B) Both Q344X and WT larvae respond a photopic light offset stimulus. The light-off VMR of WT (black trace) and Q344X (red trace) larvae at 880 lux. The light was turned off at Time = 0. Each trace shows the average larval displacement of 3 technical replicates with 48 larvae per replicate. The corresponding color ribbon indicates ± 1 s.e.m. (C) Q344X and WT larvae do not show a VMR to a scotopic light onset stimulus. The scotopic light-on VMR of WT (black trace) and Q344X (red trace) larvae at 0.01 lux. The light was turned on at Time = 0. Each trace shows the average larval displacement of 18 biological replicates with 48 larvae per condition per replicate. The corresponding color ribbon indicates ± 1 s.e.m. (D) Boxplot distribution of the average larval displacement of WT and Q344X larvae one second after scotopic light onset. The average displacement of WT larvae (*µ* ± s.e.m. ): 0.129 ± 0.013 cm, N = 18) was not significantly different from Q344X larvae (0.130 ± 0.013 cm, N = 18) (Welch’s Two Sample t-test, T = 0.08, df = 33.8, p-value = 0.93).

